# Quantum chemistry reveals the thermodynamic principles of redox biochemistry

**DOI:** 10.1101/245365

**Authors:** Adrian Jinich, Avi Flamholz, Haniu Ren, Sung-Jin Kim, Benjamin Sanchez-Lengeling, Charles A. R. Cotton, Elad Noor, Alán Aspuru-Guzik, Arren Bar-Even

**Author notes:** **Corresponding Author:** Prof. Arren Bar-Even, Max Planck Institute of Molecular Plant Physiology, Am Mühlenberg 1, 14476 Potsdam-Golm, Germany.

## Abstract

Thermodynamics dictates the structure and function of metabolism. Redox reactions drive cellular energy and material flow. Hence, accurately quantifying the thermodynamics of redox reactions should reveal key design principles that shape cellular metabolism. However, only a limited number of redox potentials have been measured experimentally, and mostly with inconsistent, poorly-reported experimental setups. Here, we develop a quantum chemistry approach for the calculation of redox potentials of biochemical reactions. We demonstrate that our method predicts experimentally measured potentials with unparalleled accuracy. We calculate the reduction potentials of all redox pairs that can be generated from biochemically relevant compounds and highlight fundamental thermodynamic trends that define cellular redox biochemistry. We further use the calculated potentials to address the question of why NAD/NADP are used as the primary cellular electron carriers, demonstrating how their physiological redox range specifically fits the reactions of central metabolism and minimizes the concentration of reactive carbonyls. The use of quantum chemistry tools, as demonstrated in this study, can revolutionize our understanding of key biochemical phenomena by enabling fast and accurate calculation of large datasets of thermodynamic values.

## Introduction

In order to understand life we need to understand the forces that support and constrain it. Thermodynamics provides the fundamental constraints that shape metabolism ^1–4^. Redox reactions constitute the primary metabolic pillars that support life. Life itself can be viewed as an electron transport process that conserves and dissipates energy in order to generate and maintain a heritable local order ^5^. Indeed, almost 40% of all known metabolic reactions are redox reactions ^6,7^. Redox biochemistry has shaped the study of diverse fields in biology, including origins-of-life ^8^, circadian clocks ^9^, carbon-fixation ^10^, cellular aging ^11^, and host-pathogen interactions ^12^. Previous work has demonstrated that a quantitative understanding of the thermodynamic parameters governing redox reactions reveals design principles of metabolic pathways. For example, the unfavorable nature of carboxyl reduction and carboxylation explains to a large degree the ATP investment required to support carbon fixation ^1^.

Developing a deep understanding of redox biochemistry requires a comprehensive and accurate set of reduction potentials values covering a broad range of reaction types. However, only ∼100 reduction potentials can be inferred from experimental data, and these suffer from inconsistencies in experimental setup and conditions. Alternatively, group contribution methods (GCM) can be used to predict a large set of Gibbs energies of formation and reduction potentials ^13^. However, the accuracy of this approach is limited, as GCM do not account for interactions between functional groups within a single molecule and GCM predictions are limited to metabolites with functional groups spanned by the model and experimental data.

Here, we develop quantum chemistry tools for the accurate prediction of the reduction potentials of a large set of redox pairs. Unlike GCM, whose smallest district unit is a functional group, quantum chemistry directly relates to the atomic and electronic configuration of a molecule, enabling *ab initio* prediction of molecular energetics. We show that the quantum chemical method can predict experimentally derived reduction potentials with considerably higher accuracy than GCM when calibrated with only two parameters. We use this method to estimate the reduction potentials of all possible redox pairs that can be generated from the KEGG database of biochemical compounds ^6,7^. This enables us to decipher general trends between and within groups of oxidoreductase reactions, which highlight design principles encoded in cellular metabolism. We specifically focus on explaining the central role of NAD(P) as electron carrier from the perspective of the redox reactions it supports and the role it plays in lowering the concentration of reactive carbonyls.

## Results

### Quantum chemical predictions of biochemical redox potentials

To facilitate our analysis we divided redox reactions into several generalized oxidoreductase groups which together cover the vast majority of redox transformations within cellular metabolism (Figure 1A): (G1) reduction of an unmodified carboxylic acid (-COO) or an activated carboxylic acid – i.e., phosphoanhydride (-COOPO_3_) or thioester (-COS-CoA) – to a carbonyl (-C=O); (G2) reduction of a carbonyl to a hydroxycarbon (-COH, i.e., alcohol); (G3) reduction of a carbonyl to an amine (-CNH _3_); and (G4) reduction of a hydroxycarbon to a hydrocarbon (-C-C-), which usually occurs via an ethylene intermediate (-C=C-).

**Figure 1.**
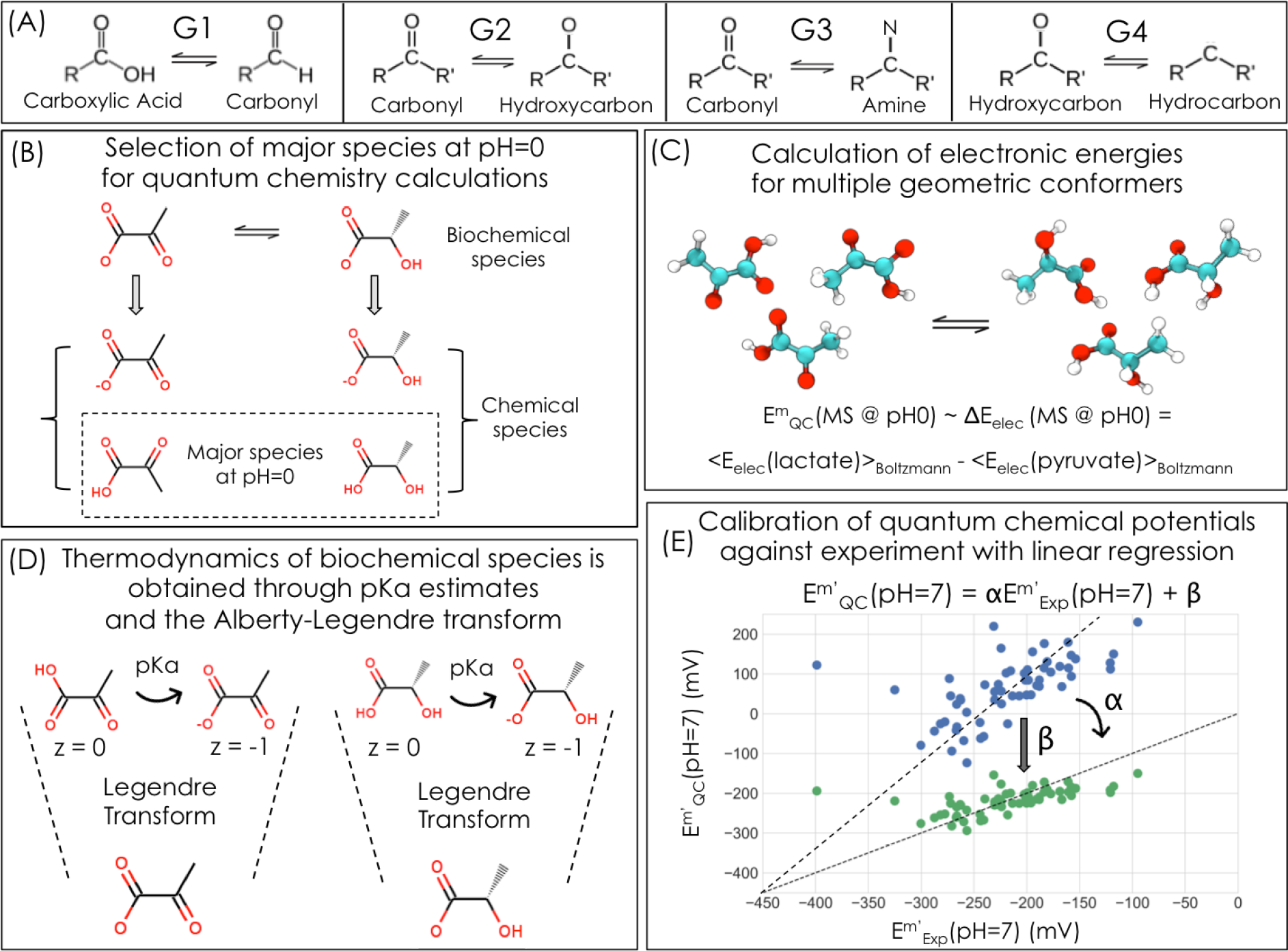
Our study is based on predicting biochemical standard redox potentials using a calibrated quantum chemistry strategy. A) The four different redox reaction categories considered here are reduction of a carboxylic acid to a carbonyl (G1), reduction of a carbonyl to a hydroxycarbon (G2), or an amine (G3), and reduction of a hydroxycarbon to a hydrocarbon (G4). B) For each redox reaction of interest, such as reduction of pyruvate to lactate, we select the most abundant protonation state at acidic pH (pH=0) for quantum chemical simulation. C) We estimate the chemical redox potential as the difference between Boltzmann-averaged electronic energies of geometric conformers of products and substrates. D) In order to convert chemical redox potentials to biochemical potentials at pH=7, we use cheminformatic pKa estimates and the Alberty-Legendre Transform (Supplementary Information). (E) Finally, we use available experimental values to calibrate redox potentials using linear regression.

We developed a quantum chemistry method for predicting the standard transformed redox potential of biochemical redox reactions. We explored a range of different model chemistries and found that a DFT (density functional theory) approach that uses the double-hybrid functional B2PLYP^14,15^ gave the highest prediction accuracy (see Methods for detailed model chemistry description; other model chemistries also gave high accuracy as discussed in the Supplementary Information and Figure S1). As each biochemical compound represents an ensemble of different chemical species – each at a different protonation state ^16^ – we applied the following pipeline to predict E’^o^ (Figure 1, see also Methods): (i) a quantum chemical simulation was used to obtain the electronic energies of the most abundant chemical species at pH 0; (ii) we then calculated the difference in electronic energies between the product and substrate of a redox pair at pH 0, thus obtaining estimates of the standard redox potential, E^o^; (iii) next, we employed empirical pKa estimates to calculate the energetics of the deprotonated chemical species and used the extended Debye-Huckel equation and the Alberty-Legendre transform ^16^ to convert E^o^ to the standard transformed redox potential E’^m^ at pH=7 and ionic strength I = 0.25 M (as recommended ^17^), where reactant concentrations are standardized to 1 mM to better approximate the physiological concentrations of metabolites ^1,18^. Finally, (iv) to correct for systematic errors, the predicted E’^m^ values, of each oxidoreductase group, were calibrated separately by linear regression (two-parameter calibration) against the set of experimentally measured potentials. We note that the two-parameter calibration is needed mainly since we ignore vibrational enthalpies and entropies of the compounds (Supplementary Information).

As exemplified in Figure 2A-B, the calibration by linear regression significantly improves the accuracy of our quantum chemistry predictions. As shown in Table 1, the predictions of quantum chemistry have a lower mean absolute error (MAE) than those of GCM for all reaction categories. (GCM has a higher Pearson correlation coefficient for category G1, but this is an artifact introduced by a single outlier value, Figure S3). The improved accuracy is especially noteworthy as our quantum chemical approach derives reduction potentials from first principles and requires only two calibration parameters per oxidoreductase group, as compared to GCM which uses 5–13 parameters while achieving lower prediction accuracy (Table 1). Therefore, our quantum chemistry approach can be extended to predict reduction potentials for a wide domain of redox reactions since it does not depend as heavily on empirical measurements.

**Figure 2.**
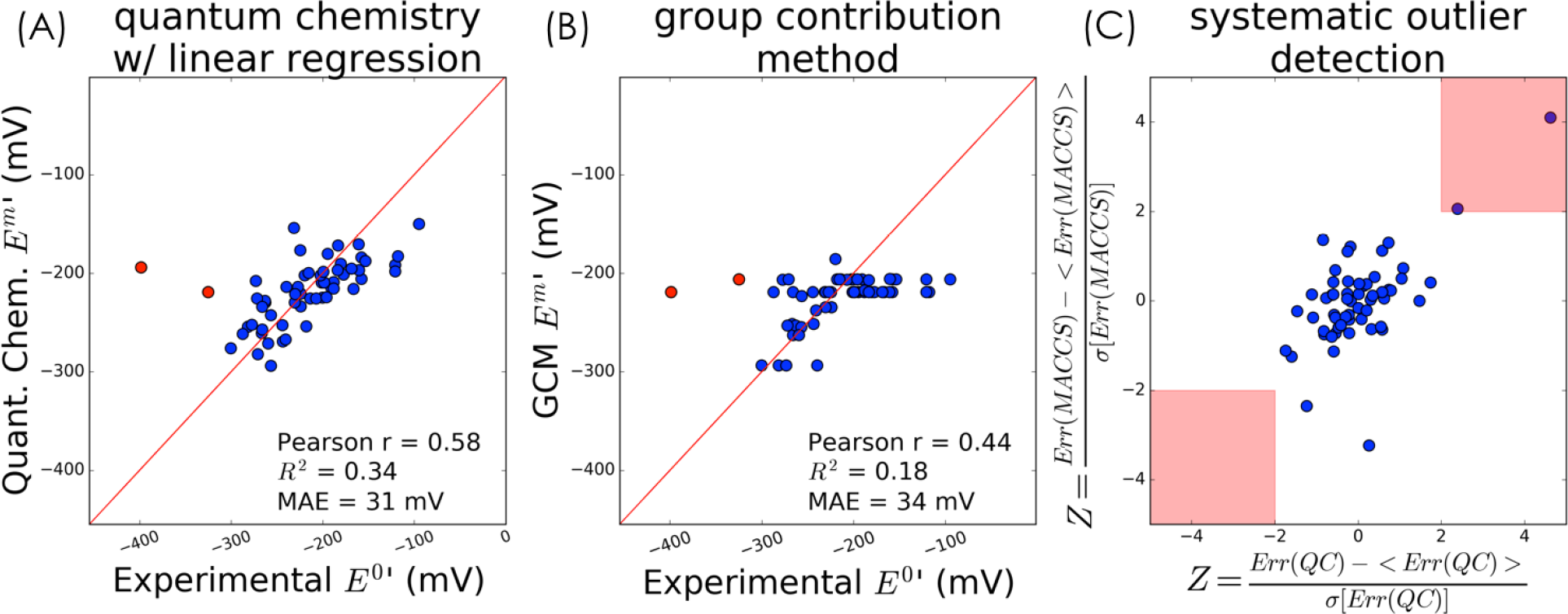
Prediction of biochemical redox potentials of carbonyl to hydroxycarbon reactions (category G2). (A) Calibration of quantum chemical estimates through linear regression (2-parameters per reaction category) significantly improves prediction accuracy (see also Figure S2). Data shown corresponds to reactions where carbonyl groups (ketones or aldehydes) are reduced to hydroxyls (G2). Quantum chemical predictions were performed using the double-hybrid DFT functional B2PLYP, the DefBas-2 basis set, COSMO implicit solvent, and D3 dispersion correction (Supplementary Information). Points in red correspond to reactions that consistently appear as outliers across modeling approaches: the indolepyruvate reduction to indolelactate and succinate semialdehyde reduction to 4-hydroxybutanoate (B) Prediction accuracy of Group Contribution Method (10 parameters for the G2 category) (C) Scatter plot of normalized prediction errors (z-scores) of G2 reactions for molecular fingerprints and quantum chemistry. The indolelactate dehydrogenase (EC 1.1.1.110) and the succinate semialdehyde reductase (EC 1.1.1.61) reactions have potentially erroneous experimental values.

**Table 1.**
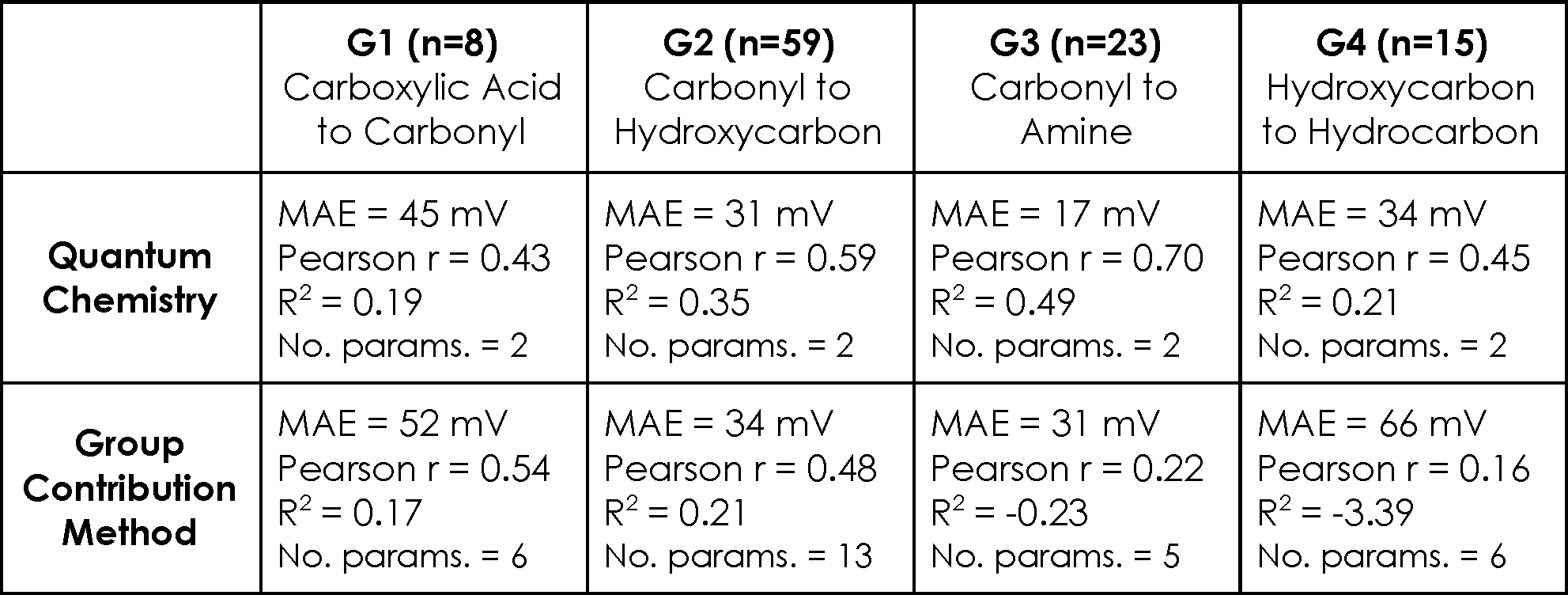
Prediction accuracy of the quantum chemistry and group contribution method modeling approaches. The number of available experimental values for each reaction category is indicated in parentheses. MAE = Mean Absolute Error; R^2^ = coefficient of determination. Note that for the G1 category, quantum chemistry has a lower MAE, but GCM has higher values of Pearson r.

|  | **G1 (n=8)**<br>Carboxylic Acid<br>to Carbonyl | **G2 (n=59)**<br>Carbonyl to<br>Hydroxycarbon | **G3 (n=23)**<br>Carbonyl to<br>Amine | **G4 (n=15)**<br>Hydroxycarbon<br>to Hydrocarbon |
| --- | --- | --- | --- | --- |
| **Quantum**<br>**Chemistry** | MAE = 45 mV<br>Pearson r = 0.43<br>$R^2 = 0.19$<br>No. params. = 2 | MAE = 31 mV<br>Pearson r = 0.59<br>$R^2 = 0.35$<br>No. params. = 2 | MAE = 17 mV<br>Pearson r = 0.70<br>$R^2 = 0.49$<br>No. params. = 2 | MAE = 34 mV<br>Pearson r = 0.45<br>$R^2 = 0.21$<br>No. params. = 2 |
| **Group**<br>**Contribution**<br>**Method** | MAE = 52 mV<br>Pearson r = 0.54<br>$R^2 = 0.17$<br>No. params. = 6 | MAE = 34 mV<br>Pearson r = 0.48<br>$R^2 = 0.21$<br>No. params. = 13 | MAE = 31 mV<br>Pearson r = 0.22<br>$R^2 = -0.23$<br>No. params. = 5 | MAE = 66 mV<br>Pearson r = 0.16<br>$R^2 = -3.39$<br>No. params. = 6 |

### Systematic detection of potentially erroneous experimental values

Inconsistencies between our predictions and experimental measurements can be used to identify erroneous experimental values. However, as a such discrepancy might stem from false prediction, we used an independent method for prediction of redox potentials. We reasoned that consistent deviation from two very different prediction approaches should be regarded as indicative of experimental error. The second prediction approach we used is based on reaction fingerprints ^19^, where the structure of the reactants involved is encoded as a binary vector (166 parameters without regularization, Supplementary Information). These binary vectors are then used as variables in a regularized regression to correlate structure against a physicochemical property of interest, such as redox potential ^19,20^. This approach is similar to the group contribution method (GCM) in that it is based on a structural decomposition of compounds; however, unlike GCM, fingerprints encode a detailed structural representation of the compounds.

To detect potentially erroneous experimental measurements, we focused on redox potentials of category G2 (carbonyl to hydroxycarbon reduction) as we have abundant experimental information for this oxidoreductase group (see Figure S4 for results with the other categories). As shown in Figure 2C, we normalized the prediction errors by computing their associated z-scores (indicating how many standard deviations a prediction error is from the mean error across all reactions). Two redox reactions stand out as having significantly different experimental and predicted values for both methods (Z>2): indolepyruvate reduction to indolelactate (indolelactate dehydrogenase, EC 1.1.1.110) and succinate semialdehyde reduction to 4-hydroxybutanoate (succinate semialdehyde reductase, 1.1.1.61).

We suggest an explanation for the observed deviation of the first reaction: in the experimental study, the K’_eq_ of indolelactate dehydrogenase was measured using absorbance at 340 nm as an indicator of the concentration of NADH ^21^. However, since indolic compounds also have strong absorption at 340 nm ^22^, this method probably resulted in an overestimation of the concentration of NADH, and thus an underestimation of K’_eq_. Indeed, the experimentally derived E’^m^ is considerably lower (-400 mV) than the predicted one (-190 mV, via quantum chemistry). With regards to the second reaction, succinate semialdehyde reductase, we note that re-measuring its redox potential is of considerable significance as it plays a central role both in carbon fixation – e.g., the 3-hydroxypropionate-4-hydroxybutyrate cycle and the dicarboxylate-4-hydroxybutyrate cycle ^10^ – as well as in production of key commodities – e.g., biosynthesis of 1,4-butanediol ^23^.

### Comprehensive prediction and analysis of reduction potentials

We used the calibrated quantum chemistry to predict redox potential for a database of natural and non-natural redox reactions. We generated this dataset by identifying pairs of metabolites from KEGG ^6,7^ that fit the chemical transformations associated with each of the four different oxidoreductase groups (Methods). We considered only compounds with fewer than 7 carbon atoms, thus generating a dataset consisting of 652 reactions: 83 reductions of category G1; 205 reductions of category G2; 104 reductions of category G3; and 260 reductions of category G4 (Supplementary Dataset 1). Some of these redox pairs are known to participate in enzyme-catalyzed reactions while others are hypothetical transformations that could be performed by engineered enzymes.

Figure 3A shows the distribution of all predicted redox potentials at pH = 7, I = 0.25 M and reactant concentrations of 1 mM, i.e., E’^m^ ^13,24^. Figure 3 demonstrates that the value of E’^m^ is directly related to the oxidation state of the functional group being reduced. The general trend is that “the rich get richer” ^1,25,26^: more reduced functional groups have a greater tendency to accept electrons, i.e., have higher reduction potentials. Specifically, the reduction potential of hydroxycarbons (G4, < *E’^m^>* = − 15 *mV*) is higher than that of carbonyls (<*E’^m^>* = − 225 *mV* for both G2 and G3) and the reduction potential of carbonyls is higher than that of un-activated carboxylic acids (G3, < *E’^m^>* = − 550 *mV*). Categories G2 and G3 (reduction of carbonyls to hydroxycarbons or amines,respectively) have very similar potentials because the oxidation state of the functional groups involved is identical^1^. For category G1, activation of carboxylic acids significantly increases their reduction potential (orange line in Figure 3) as the energy released by the hydrolysis of the phosphoanhydride or thioester (∼50 kJ/mol) activates the reduction: Δ*E* = 50/(*nF*) ≃ 250 *mV* (n being the number of electrons, F the Faraday constant).

**Figure 3.**
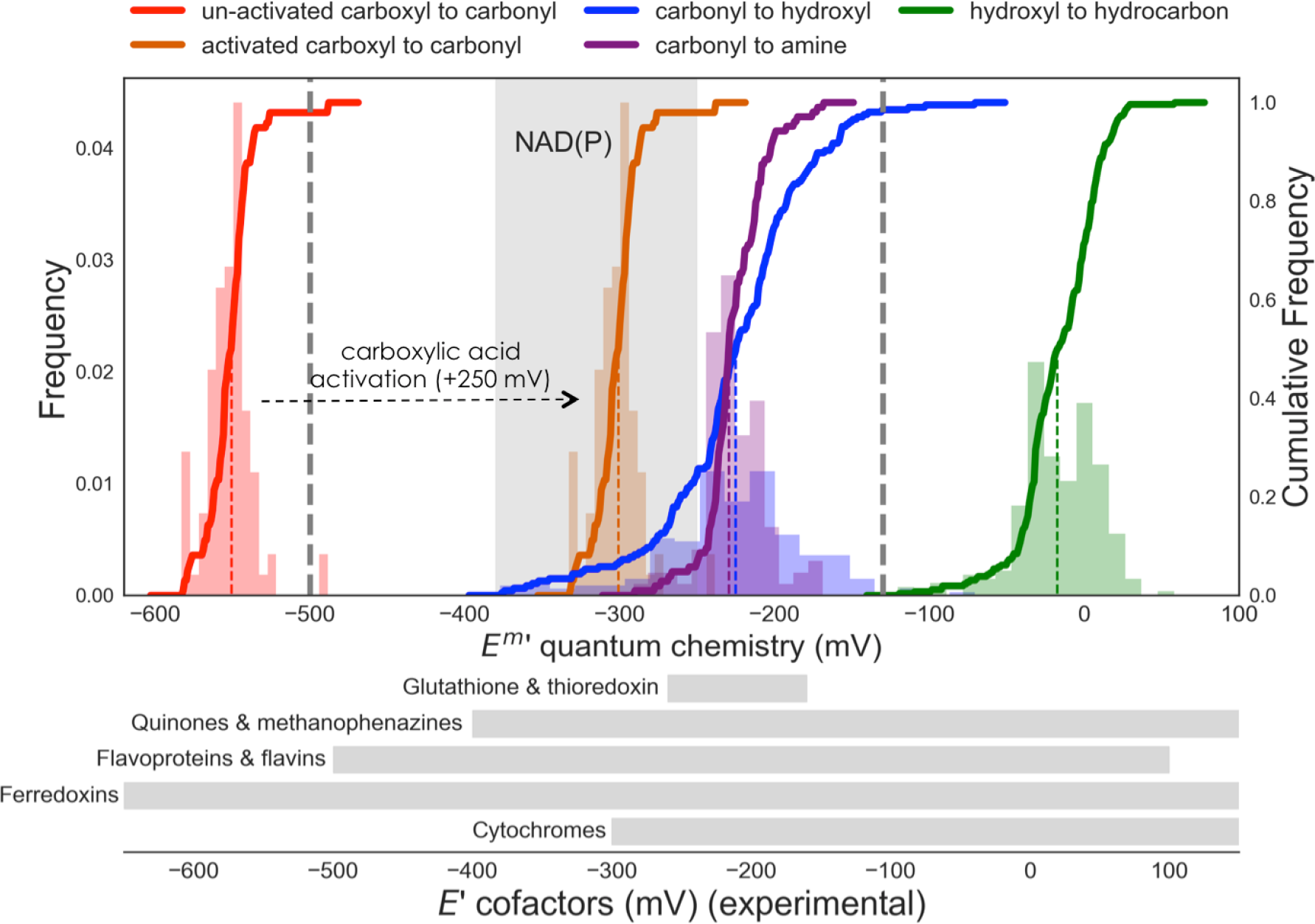
Distributions of predicted standard transformed redox potentials at pH = 7 and I = 0.25 for a dataset of 650 natural and non-natural reactions. The average reduction potentials for each reaction category are: un-activated carboxylic acid to carbonyl (G1: <*E’^m^* > = − 550 *mV*), activated carboxylic acid to carbonyl (activated G1: < *E’^m^* > = − 300 *mV*), carbonyl to hydroxycarbon (G2: < *E’^m^* > = − 225 *mV*), carbonyl to amine (G3: < *E’^m^* > = − 225 *mV*), and hydroxycarbon to hydrocarbon (G4: < *E’^m^* > = − 15 *mV*) Both histograms and cumulative distributions (bold lines, right y-axis) are shown. The distributions for unactivated and activated carboxylic acid to carbonyl reductions (red and purple) are the same, but shifted by +250 mV. Dashed colored lines show the median redox potential for each reaction category. Grey shaded regions corresponds to the range of NAD(P) redox potential, while dashed grey lines delimit the region of reversible oxidation/reduction by NAD(P)/NAD(P)H. Ranges of reduction potentials for different alternative cofactors are shown as grey rectangles (Table S2).

The quantum chemical predictions further enable us to explore detailed structure-energy relationships within each of the general oxidoreductase groups. To exemplify this we focus on the G2 category, as shown in Figure 4. While we find no significant difference between the average E’^m^ of aldehydes and ketones, we can clearly see that the identity of functional groups adjacent to the carbonyl has a significant effect on E’^m^, as expected. Alpha ketoacids and dicarbonyls have a significantly higher E’^m^ than alpha hydroxy-carbonyls (Δ <*E’^m^* ≃ = 20 *mV*, *p* < 0.005)and carbonyls adjacent to hydrocarbons (Δ <*E’^m^* ≃ = 35 *mV*, *p* < 0.005). Carbonyls next to double bonds or aromatic rings have a significantly lower E’^m^ values than alpha hydroxy-carbonyls and carbonyls that are next to hydrocarbons (Δ <*E’^m^* > ≃ −50 *mV*, and Δ <*E’^m^* > ≃ − 40 *mV*, *p* < 0.0001), respectively, *p* < 0.0001 *p* < 0.0001). Lactones (cyclic esters), have redox potentials that are significantly lower than any other subgroup within the G2 category. As another validation of the predicted potentials, we found that the reduction potentials of open-chain sugars are significantly higher than those of closed-ring sugars that undergo ring opening upon reduction, where Δ <*E’^m^* > ≃ − 60*mV, p* < 0.0001. This is consistent with the thermodynamics of closed-ring sugar conformations, e.g., the K_eq_ of arabinose ring opening is ∼350 (25), which translates to Δ*E* = *RTln*(350)/(*nF*) ≃ 75 *mV*, close to the observed average potential difference between the subgroups (R is the gas constant, and T the temperature).

**Figure 4.**
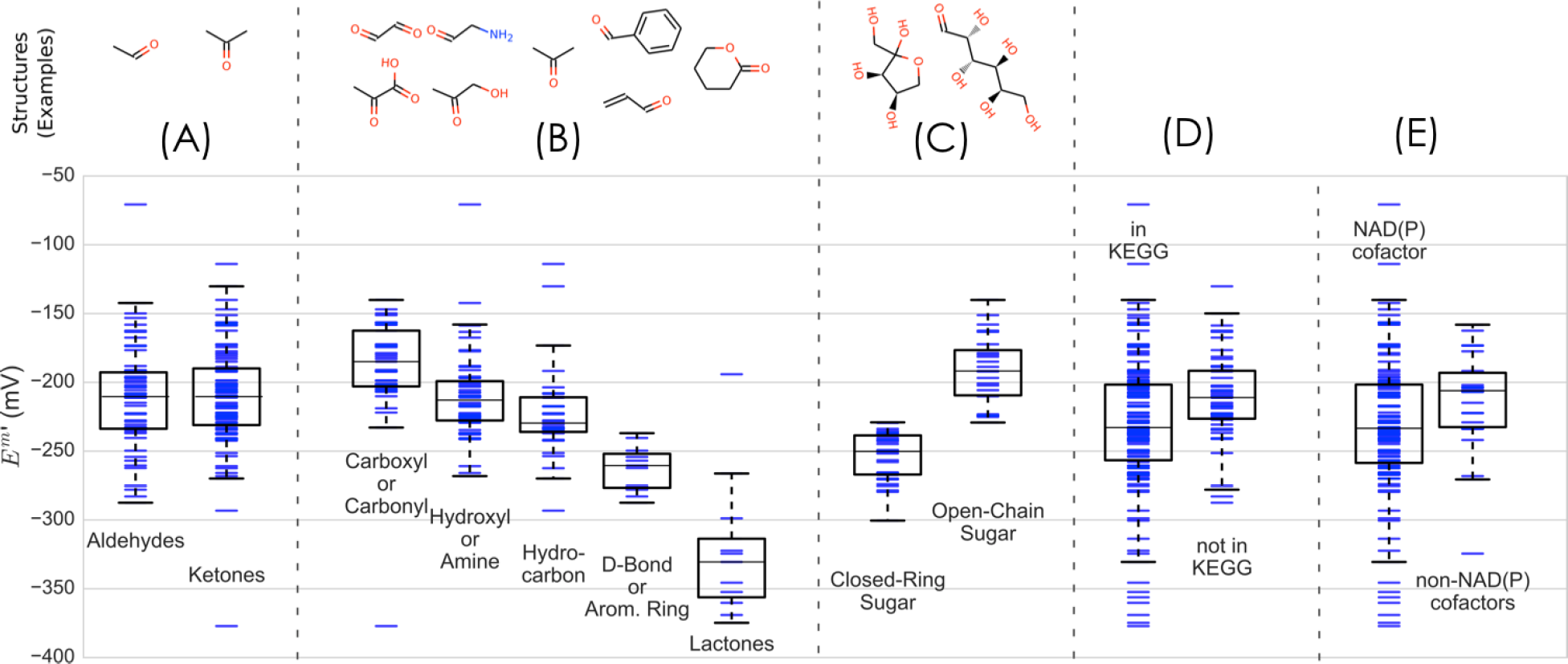
Structure-energy relationships in the standard transformed redox potentials for reactions in the G2 category (carbonyl to hydroxycarbon reductions). Five different modes of analysis were considered. (A) Aldehydes vs. ketones (non-statistically significant Δ *E*’*m* >); (B) nearest-neighbor functional group (all subgroups have statistically significant Δ *E*’*m* >, *p* < 0.005); except hydroxyl/amine and hydrocarbon) (C) closed-ring sugar reduction to open-chain vs. open-chain sugar reduction to open-chain (statistically significant Δ *E*’*m* >, *p* < 10^−5^), (D) natural reactions appearing in KEGG vs. non-natural reactions (statistically significant Δ *E*’*m* >, *p* < 0.005) (E) natural reactions that only use NAD(P) as redox cofactor vs. those that use alternative cofactors (cytochromes, FAD, O_2_, or quinones) (non-statistically significant Δ < *E*’*m* >, *p* < 0.003).

### On the biochemical logic of the universal reliance of NAD(P)

While myriad natural electron carriers are known to support cellular redox reactions, NAD(P) has the prime role in almost all organisms, participating in most (>50%) known redox reactions ^6,7^. The standard redox potential of NAD(P) is ∼−330 mV (pH=7, I=0.25), but as [*NADPH*]/[*NADP*] can be higher than 50 and [*NADPH*]/[*NADP*] can be lower than 1/500, the physiological range of the NAD(P) reduction potential is between −380 mV and −250 mV (16, 26–30) ^18,27–31^. Most cellular redox reactions are therefore constrained to a limited reduction potential range determined by the physicochemical properties and physiological concentrations of NAD(P). By examining the fundamental trends of redox potentials of the different oxidoreductase groups we will show that NAD(P) is well-matched to the redox transformations most commonly found in cellular metabolism.

Figure 3A demonstrates that the reduction potentials of activated acids (activated G1) and carbonyls (to hydroxyls and amines, G2 and G3) are very similar, such that NAD(P) can support both the oxidation and reduction of nearly all redox couples in these classes. Although the distributions associated with these redox reactions are not entirely contained in the NAD(P) reduction potential range (marked in grey), the reduction potential of a redox pair can be altered by modulating the concentrations of the oxidized and reduced species. As the concentrations of metabolites usually lie between 1 μM and 10 mM (1, 4, 16, 31) ^1,4,18,32^, the reduction potential of a redox pair can be offset from its standard value by up to ±*RT ln*(10^4^)/(*nF*) ≈ ±120 *mV* (assuming two electrons are transferred). Therefore, NAD(P) can support reversible redox reactions of compound pairs with E’^m^ as −380 −120 = −500*mV* low as and as high as −250+120 = − 130 *mV* (indicated by the dashed lines in Figure 3), a range that encompasses almost all activated acids (activated G1) and carbonyls (G2 and G3 reactions). Outside this range, however, NAD(P)(H) can only be used in one direction of the redox transformation – either oxidation or reduction, but not both. Figure 3 shows that NAD(P)H can support irreversible reductions of hydroxycarbons to hydrocarbons and NAD(P) supports irreversible oxidation of carbonyls to carboxylic acids.

Next, we focus on a small set of redox reactions found in the extended central metabolic network that is shared by almost all organisms: (i) The TCA cycle, operating in the oxidative or reductive direction ^33,34^, as a cycle or as a fork ^35^, being complete or incomplete ^35^, or with some local bypasses (e.g., ^36^); (ii) glycolysis and gluconeogenesis, whether via the EMP or ED pathway ^37^, having fully, semi or non-phosphorylated intermediates ^38^; (iii) the pentose phosphate cycle, working in the oxidative, reductive or neutral direction; and (iv) biosynthesis of amino-acids, nucleobases and fatty acids. As schematically shown in Figure 5, and listed in Supplementary Dataset S2, the ≈ 60 redox reactions that participate in the extended central metabolism almost exclusively belong to one of the following groups: (i) reduction of an activated carboxylic acid to a carbonyl or the reverse reaction oxidizing the carbonyl (9 reactions, G1); (ii) reduction of a carbonyl to a hydroxycarbon or its reverse oxidation (20 reactions, G2); (iii) reduction of a carbonyl to an amine or its reverse oxidation (18 reactions, G3); (iv) irreversible oxidation of carbonyls to un-activated carboxylic acids (5 reactions, G1 in the direction of oxidation); and (v) irreversible reduction of hydroxycarbon to hydrocarbons (4 reactions, G4). Only two central metabolic reactions (marked in magenta background in Figure 5) oxidize hydrocarbons to hydroxycarbons (G4, in the direction of oxidation) and require a reduction potential higher than that of NAD(P): oxidation of succinate to fumarate and oxidation of dihydroorotate to orotate^2^. Similarly, the extended central metabolic network does not demand the low reduction potential required for the reduction of un-activated carboxylic acids (G1).

**Figure 5.**
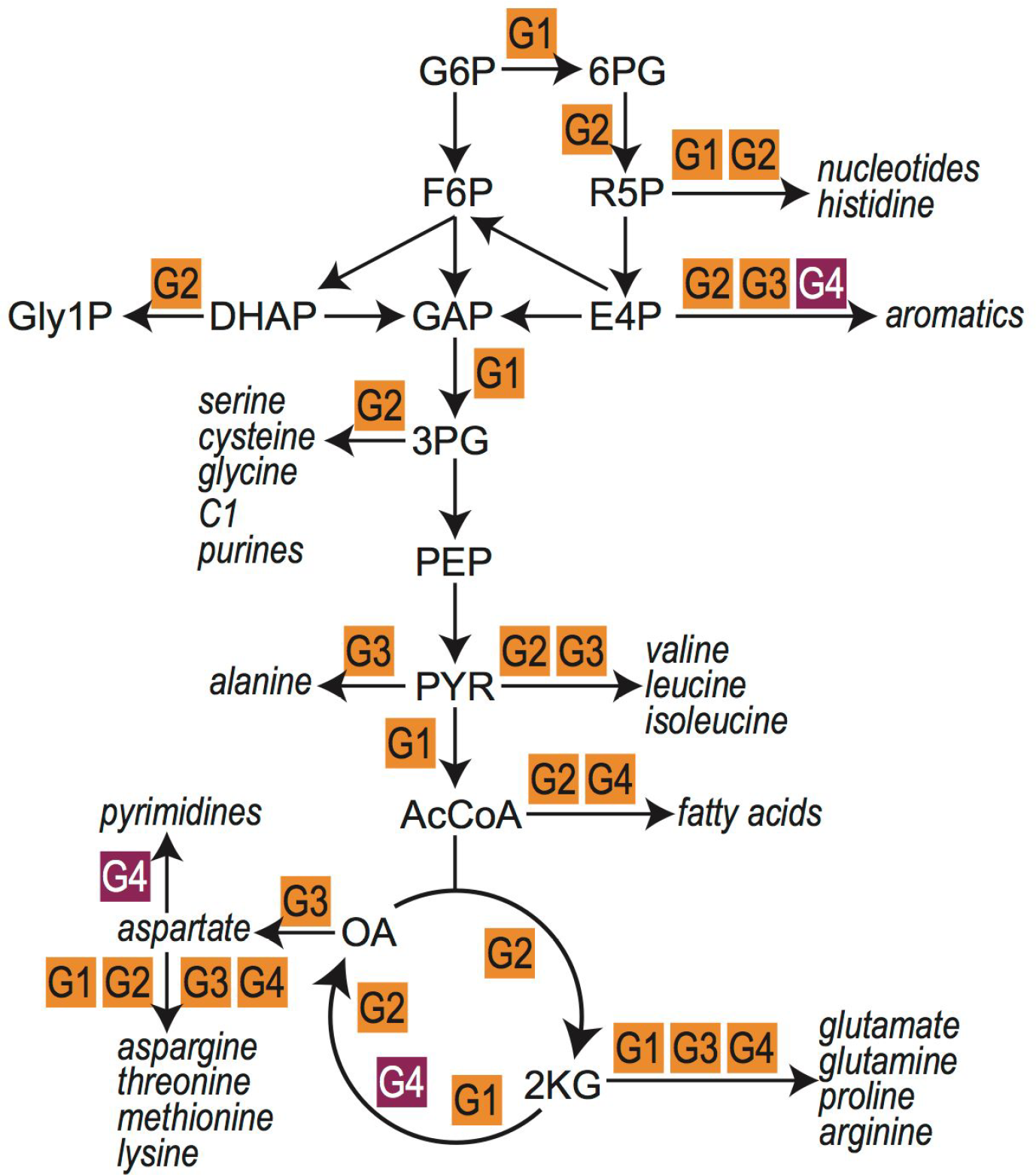
A schematic localization of the types of oxidoreductase reactions (oxidoreductase groups 1 to 4) within the extended central metabolic network. We highlights reactions (purple) where a hydrocarbon is oxidized to a hydroxycarbon (G4 reactions, in the direction of oxidation) which generally cannot be sustained by NAD(P) as redox cofactor. See Supplementary Dataset 2 for full set of redox reactions in extended central metabolic network. G6P, Glucose-6-phosphate; F6P, Fructse-6-phosphate; DHAP, Dihydroxyacetone phosphate; GAP, Glyceraldehyde 3-phosphate; Gly1P; Glycerol 1-phosphate; 6PG, 6-Phosphogluconolactone; R5P, Ribulose 5-phosphate; E4P, Erythrose 4-phosphate; 3PG, 3-Phosphoglycerate; PEP, Phosphoenolpyruvate; PYR, Pyruvate; AcCoA, Acetyl coenzyme A; 2KG, 2-Ketoglutaric acid; OA, Oxaloacetate.

The reduction potential range associated with NAD(P) therefore perfectly matches the vast majority of reversible redox reactions in extended central metabolism – i.e., reduction of activated carboxylic acids and reduction of carbonyls (orange, purple and blue distributions in Figure 3) – and can also support the common irreversible redox transformations of extended central metabolism – i.e., reduction of hydroxycarbons and oxidation of carbonyls to un-activated carboxylic acids (green and red distributions in Figure 3). Cells typically rely on secondary redox carriers like quinones and ferredoxins (Figure 3, Table S3), to support less common reactions, i.e., oxidation of hydrocarbons and reduction of un-activated carboxylic acids.

*Why is the reduction potential of NAD(P) lower than the *E’^m^* of most carbonyls* (Figure 3)? As biosynthesis of an NAD(P) derivative with higher reduction potential presents no major challenge (28), *why does this lower potential persist*? We suggest that this redox offset plays an important role in reducing the concentrations of cellular carbonyls by making their reduction to hydroxycarbons favorable. It is well known that carbonyls are reactive towards macromolecules, as they spontaneously cross-link proteins, inactivate enzymes and mutagenize DNA ^39,40^. As the reduction potential of NAD(P) is lower than most carbonyls, the redox reactions in category G2 (or G3) prefer the direction of reduction, thus ensuring that carbonyls are kept at lower concentrations than their corresponding hydroxycarbons (or amines). Assuming a value of E’ = −330 mV for NAD(P) and taking the average E’^m^ of the G2 reactions (<*E’^m^* > = − 25 *mV*) results in an estimated equilibrium concentration ratio, [*hydroxycarbon*] / [*carbonyl*] *exp*(*E*’[*NAD*(*P*)] −<*E’^m^*)*nF* /*RT*) ≃ 3500 thus ensuring very low levels of the carbonyl species.

For ketoacids and open-ring sugars (which are especially reactive due to the free carbonyl) this effect is even more pronounced as both have especially high reduction potentials (Figure 4). Indeed, the reduction potential of ketoacids is so high that the reverse, oxidative reaction is usually supported by electron donors with a higher potential than NAD(P), for example, quinones, flavins, and even O_2_ (e.g., lactate oxidase, glycolate oxidase). Interestingly, the reactions of category G2 that are supported by known enzymes in the KEGG database (75% of reactions in this category) have significantly lower E’^m^ than the remaining reactions, which are not known to be catalyzed by natural enzymes (Δ < *E’^m^* > ≃ 20 *mV, p* < 0.005). As such, we suggest that the G2 transformations that are known to be enzyme-catalyzed are mainly those that are amenable to redox coupling with NAD(P) (Figure 4D). Within the subset of G2 transformations found in KEGG, those that use redox cofactors other than NAD(P) (such as cytochromes, FAD, O_2_, or quinones) have higher E’^m^ values (Δ <*E’^m^* > ≃ 20 *mV*, not significant *p* = 0.03) than those that use NAD(P) (Figure 4).

Finally, we note that the reduction potential of NADP and activated carboxylic acids (activated G1) overlap almost completely, such that we would not expect NAD(P) to have a strong effect on the ratio between the concentrations of carbonyls and activated acids. This is to be expected as both carbonyls and activated carboxylic acids are reactive – e.g., acetylphosphate and glycerate bisphosphate acetylates proteins spontaneously ^41^ and acyl-CoA’s S-acetylates cellular peptides non-enzymatically ^42^. As such, there is no sense in driving the accumulation of carbonyls at the expense of activated carboxylic acids or vice-versa – neither approach would ameliorate non-specific toxicity.

## Discussion

In this work, we present a novel approach for predicting the thermodynamics of biochemical redox reactions. Our approach differs radically from group contribution methods, which rely on a large set of arbitrarily-defined functional groups, assume no energetic interactions between groups, and are restricted to metabolites that are decomposable into the groups spanned by the model. In contrast, quantum chemistry directly takes into account the electronic structure of metabolites in solution. The current implementation relies on a two-parameter calibration for each oxidoreductase reaction category, which reduces computational cost by avoiding the need to calculate vibrational enthalpies and entropies (Supplementary Information). In future studies, faster computational resources will enable full *ab initio* prediction of hundreds of standard transformed redox potentials, rendering the two-parameter calibration and the use of empirical pKa values obsolete ^43^. Yet, as we have shown, the current procedure is sufficient to yield high coverage and accuracy at a reasonable computational cost.

Importantly, the quantum chemical strategy is not subject to the inconsistencies that plague experimental databases. Experimental values are measured in a wide range of different conditions, including temperature, pH, ionic strength, buffers, and electrolytes. In many cases, the exact measurement conditions are not reported, making it practically impossible to account for these factors. Thus, even if we were to gain access to more experimental data, the lack of systematically applied conditions makes such resources problematic. In contrast, quantum chemical simulations can be performed in consistent, well-defined conditions.

Systematic prediction of the thermodynamic values governing key biochemical processes can reveal hidden design principles of cellular metabolism. In particular, the enhanced resolution provided by quantum chemistry uncovers important patterns not accessible using traditional analyses. Exemplifying this, we found that the main cellular electron carrier, NAD(P), is ‘tuned’ to reduce the concentration of reactive carbonyls, thereby keeping the cellular environment more chemically stable. Yet, this protection comes at a price: the oxidation of hydroxycarbons is thermodynamically challenging and often requires the use of electron carriers with higher reduction potential. A recent study demonstrates the physiological relevance of this thermodynamic barrier: the NAD-dependent 3-phosphoglycerate dehydrogenase – the first enzyme in the serine biosynthesis route – can sustain high flux in spite of its unfavorable thermodynamics only through coupling with the favorable oxidation of 2-hydroxyglutarate ^44^.

Our analysis further supports the previous assertion that the TCA cycle has evolved in the reductive direction (32, 42)^33,45^. While all other the electron transfer reactions in the extended central metabolism belongs to oxidoreductase groups that can be supported by NAD(P)(H), oxidation of succinate – a key TCA cycle reaction – cannot be carried by this electron carrier. As the reverse reaction, i.e., fumarate reduction, can be support by NADH ^46,47^, it is reasonable to speculate that the reaction first evolved in the reductive direction, and only later was adapted to work in the oxidative direction using an alternative cofactor.

So long as sufficient experimental data is available to allow for calibration, our approach can be extended to other types of biochemical reactions. For example, understanding the thermodynamics of carboxylating and decarboxylating enzymes – the “biochemical gateways” connecting the inorganic and the organic world – could pave the way for the identification of highly efficient, thermodynamically favorable carbon fixation pathways based on non-standard but promising reaction chemistries ^48,49^. In this way, high-resolution thermodynamic analyses may provide much needed insight for the engineering of microbes to address global challenges.

## Methods

We performed quantum chemical simulations on the major species (MS) of each metabolite of interest at pH = 0, which corresponds to the most positively charged species. Running calculations on these reference protonation states yields estimates for the standard redox potential, E^o^ (major species at pH=0). Using pKa values from ChemAxon’s calculator plugin (Marvin 17.7.0, 2017, ChemAxon) and the Alberty-Legendre transform, we converted E^o^ (MS at pH=0) to the standard transformed redox potential of interest, E’^m^ (pH=7, I=0.25) ^16,17^.

### Quantum chemical geometry optimizations

For each metabolite, we generated ten initial geometric conformations using ChemAxon’s calculator plugin. Quantum chemistry calculations were performed using the Orca software package (version 3.0.3) ^50^. Geometry optimizations were carried out using DFT, with the B3LYP (Becke 1993) functional and Orca’s predefined DefBas-2 basis set (see Table S3 for detailed basis set description). The COSMO implicit solvent model ^51^ was used, with the default parameter values of epsilon = 80.4 and refrac=1.33. DFT-D3 dispersion correction ^52^ using Becke-Johnson damping ^53^ was also included. Molecular hydrogen was included in the substrate side of the half-reactions (in the direction of reduction).

### Quantum chemical electronic single point energies (SPE) and calibration against experimental values

Single point energy (SPE) calculations yield the value of the electronic energy *E_Electronic_* for each conformer at their optimized geometry. We used the optimized geometries obtained using DFT as inputs for SPE calculations. Substrate and product conformers were sampled according to a Boltzmann distribution. By taking the difference of products’ and substrates’ *E_Electronic_* values, we obtain Δ*E_Electronic_*, which we treat as directly proportional to the standard reduction potential of the major species at pH 0:

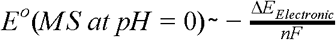

The use of Δ*E_Electronic_* to approximate the reduction potential as opposed to Δ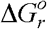 (which includes rotational and vibrational enthalpies and entropies) reduces computational cost and is motivated by the empirical observation that there is a strong correlation between Δ*E*_*Electronic*_ and 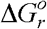 for these redox systems (Figure S5, see S1 for details). We note that we subtracted the energy of molecular hydrogen (obtained with the same SPE model chemistry) from Δ*E*_*Electronic*_ in order to get redox the potentials relative to the standard hydrogen electrode. A similar approach has been used to model redox reactions in the context of organic redox flow batteries ^54^.

For each of the redox categories, we performed linear regressions between the values of Δ*E_Electronic_* and the available experimental redox potentials. To do this, we first use cheminformatic pKa estimates and the Alberty-Legendre transform to convert both the experimental standard redox potentials and the quantum chemical predictions of E^o^(MS at pH=0) to the standard transformed redox potentials *E’^o^* (*pH* = 7, *I* =0). The calibration via linear regression was implemented using the SciKit learn Python library (http://scikit-learn.org/).

In order to optimize prediction accuracy we ran SPE calculations using a large diversity of model chemistries (Figure S1, and see SI for details). Optimizing for Pearson correlation coefficients r, we selected the following model chemistry to predict reactions without experimentally measured E’^m^: a DFT approach with the double-hybrid functional B2PLYP, the DefBas-5 Orca basis set (see Table S3 for detailed basis set description), COSMO implicit solvent, and D3 dispersion correction. To avoid overfitting, we trained the model chemistry optimization procedure on the experimental data for the G3 reaction category (carbonyl to amine reduction), and validate its accuracy on the rest of the oxidoreductase reaction categories (Table 1 and Table S2).

Although we explored a large set of DFT functionals, wavefunction methods, and basis sets, further improvements can be achieved by exploring a larger space of model chemistries, including the geometry optimization procedure, conformer generation method, as well as explicit solvation models ^55^. For example, adapting a recent highly accurate method (tested on four molecules) based on the Linear Response Approximation (LRA) to the large scale prediction of E’^m^ values would be an interesting direction ^43^.

### Predicting redox potentials with molecular fingerprints and group contribution method

We used the RDKit cheminformatics software tool, to obtain binary molecular fingerprints of each compound of interest. Because of the relatively small size of our training sets and in order to minimize overfitting, we used MACCS Key 166 fingerprints instead of other popular Morgan circular fingerprints ^56^. We concatenated each redox half-reaction substrate/product fingerprint pair into a single reaction fingerprint ^19^ and used these as input training data for regularized linear regression. We then performed an independent regularized regression for each of the four different redox reaction categories. To obtain group contribution estimates of redox potentials, we use the group matrix and the group energies of Noor *et al*. ^24^.

### Systematic detection of reactions with potentially erroneous experimental values

We design a strategy to detect reactions with potentially erroneous experimental values as listed in the NIST thermodynamics of enzyme-catalyzed reactions database (TECRDB) ^57^. We identify reactions whose predicted potential deviates from experiment by a similar amount for both the calibrated quantum chemistry and fingerprint-based modeling approaches. In order to make the errors associated to the two different modeling methods comparable, we normalize the prediction errors by computing their associated z-scores: *Z_Err_* = (*Err − μ*)/σ. We set a threshold value for the z-score of *Z* = 2, such that reactions with *Z_Err_* (*QC*) > 2 and *Z_Err_*(*MACCS*) > 2 are assigned a high likelihood of having an erroneously tabulated experimental value in NIST-TECRDB.

### Generation of comprehensive database of natural and non-natural redox reactions

To generate a database of all possible redox reactions involving natural compounds, we use a decomposition of all metabolites into functional groups as per the GCM ^24^. We find pairs of metabolites in the KEGG database with functional group vectors whose difference matches the reaction signature of any of the redox reaction categories of interest. For example, pairs of metabolites in the G1 category will have a group difference vector with a +1/−1 in the element corresponding to an carbonyl/carboxylic acid functional group respectively (see SI for details).

Using this method we succeeded in generating a rough database of redox reactions. However, additional manual and semi-automated data cleansing was required to get the final version of the database (see SI for further details). For example, use of the group difference vectors failed to account for the chirality of the metabolites, and in some instances stereochemistry was not maintained throughout the reaction. In order to solve this, we applied an additional filter, which used the conventions for assigning chirality (R/S, L/D) present in molecule names to match chirality between the substrate and product. Sugars proved to be especially problematic as those reactions did not maintain stereochemistry throughout; for these reactions, the above filtering method did not suffice, often keeping incorrect reactions such as L-Xylonate → L-Arabinose. For this, we used molecular naming conventions to eliminate the wrong reactions.

### Statistically significant differences between average E’^m^ values for distinct structural groups

We performed Welch’s unequal variance t-test to obtain the p-value for the null hypothesis that pairs of different reaction subcategories within group G2 have identical average E’^m^ values (Figure 4). Welch’s t-test is an adaptation of Student’s t-test which does not assume equal variances.

## Footnotes

1 Note that this holds for the physiological E’^m^ but not for E’^o^ because reactions in the G3 category are balanced with an ammonia molecule as a substrate, thus introducing a factor of RTln(10^−3^) when converting to the mM standard state.

2 While formally being oxidation of hydrocarbon to hydroxycarbon, the oxidations of prephenate to 4-hydroxyphenylpyruvate and of arogenate to tyrosine present a special case since they create a highly stable aromatic ring and hence have enough energy to donate their electrons directly to NAD(P).

